# Succinate can shuttle reducing power from the hypoxic retina to the O_2_-rich pigment epithelium

**DOI:** 10.1101/2020.01.17.909929

**Authors:** Celia M. Bisbach, Daniel T. Hass, Brian M. Robbings, Austin M. Rountree, Martin Sadilek, Ian R. Sweet, James B. Hurley

**Affiliations:** University of Washington

## Abstract

When O_2_ is plentiful, the mitochondrial electron transport chain uses it as a terminal electron acceptor. However, the mammalian retina thrives within a hypoxic niche in the eye. We find that mitochondria in retinas adapt to their hypoxic environment by relying on the “reverse” succinate dehydrogenase reaction, where fumarate accepts electrons instead of O_2_. Reverse succinate dehydrogenase activity produces succinate and is enhanced by down-regulation of cytochrome oxidase subunits. Retinas can export the succinate to the neighboring retinal pigment epithelium/choroid complex. There, succinate stimulates O_2_ consumption several fold and enhances synthesis and release of malate. Malate released from the pigment epithelium can be imported into the retina, where it is converted to fumarate and again used to accept electrons in the reverse succinate dehydrogenase reaction. Our findings show how a malate/succinate shuttle can sustain these two tissues by transferring reducing power from an O_2_-poor tissue (retina) to an O_2_-rich one (retinal pigment epithelium/choroid).

## Introduction

O_2_ is a substrate in one of the most important and well-known reactions of energy metabolism. Normally, it is the terminal electron acceptor in the mitochondrial electron transport chain (ETC). It has been suggested that when O_2_ is limiting, electrons from the electron transport chain can instead be passed onto fumarate. In this “reverse” succinate dehydrogenase (SDH) reaction, SDH removes electrons from the ETC to reduce fumarate to succinate, bypassing the need for O_2_ (Chouchani et al., 2014; Hochachka et al., 1975). Succinate accumulates in muscle, heart, kidney, liver, brain and blood during hypoxia (Cascarano et al., 1976; Chouchani et al., 2014; Hochachka et al., 1975). However, the degree to which the reverse SDH reaction contributes to succinate produced during hypoxia is debated, and the role of succinate in tissues that are in chronically hypoxic niches is largely unexplored (Chinopoulos, 2019; Chouchani et al., 2014; Zhang et al., 2018).

The unique architecture of the vertebrate eye places the retina in a chronically hypoxic niche. The choroidal vasculature inside the sclera of the eye is the main source of O_2_ for the outer retina. A collagenous layer and a monolayer of cells, the retinal pigment epithelium (RPE), form a barrier that selectively allows gases and nutrients to flow to the outer retina from the choroid. This results in abundant O_2_ in the RPE and a steeply declining gradient of O_2_ through the outer retina. The extent of hypoxia in the outer retina varies across species, but pO_2_ in retinas can reach as low as ∼5 mm Hg in mouse and 0 mm Hg in larger mammals (Linsenmeier and Zhang, 2017; Yu and Cringle, 2006).

To better understand the physiological consequences of this disparity in O_2_ tension, we asked how the retina and RPE have adapted to their O_2_ environments in an eye. Retinas already are known to be very glycolytic (Chinchore et al., 2017; Kanow et al., 2017; Krebs, 1927; Winkler, 1981). We discovered that retinas also adapt to hypoxia by reducing fumarate to succinate and exporting the succinate. This is the major pathway for succinate production in the retina. We find that retinas favor fumarate as an electron acceptor because the normal hypoxic state of the retina causes it to down-regulate subunits of mitochondrial Complex IV, limiting its ability to use O_2_ to accept electrons.

These observations about retinal metabolism prompted us to explore the role of succinate in the overall metabolic ecosystem of the eye. The RPE relies on its mitochondria to oxidize diverse fuels including lactate, fatty acids, glutamine and proline, and some of these fuels can be supplied to the RPE by the retina (Adijanto et al., 2014; Du et al., 2016; Kanow et al., 2017; Reyes-Reveles et al., 2017; Yam et al., 2019). In this report we show that the RPE/choroid complex has an extraordinary capacity to oxidize succinate. When fueled with succinate, the RPE/choroid complex releases malate, which can be converted back into succinate in retinas by reverse SDH activity.

Based on these findings, we propose that succinate acts as a shuttle to transfer unused reducing power from the hypoxic retina to the RPE. The RPE is an O_2_-rich tissue that is well-situated to use the residual reducing power in succinate to reduce O_2_ to H_2_O. This is consistent with a metabolic cycle in the eye in which a succinate/malate shuttle transfers electrons from the hypoxic retina to O_2_ in the RPE/choroid complex.

## RESULTS

### Retinas release succinate which can fuel O_2_ consumption in eyecups

Retinas in an eye are in a chronically hypoxic environment (Linsenmeier and Zhang, 2017; Yu and Cringle, 2006). Exposure to hypoxia can induce tissues to release succinate, so we asked if freshly isolated mouse retinas export succinate (Cascarano et al., 1976; Chouchani et al., 2014; Hochachka et al., 1975). We incubated retinas from 4-6 month old C57BL/6J mice in 5 mM ^12^C-glucose at atmospheric (21%) O_2_ and 5% CO_2_ and then measured the concentrations of metabolites exported into the medium after 30, 60, and 90 minutes (**Figure 1A**). Succinate is the most abundant TCA cycle metabolite released by retinas under these conditions.

**FIGURE 1:**
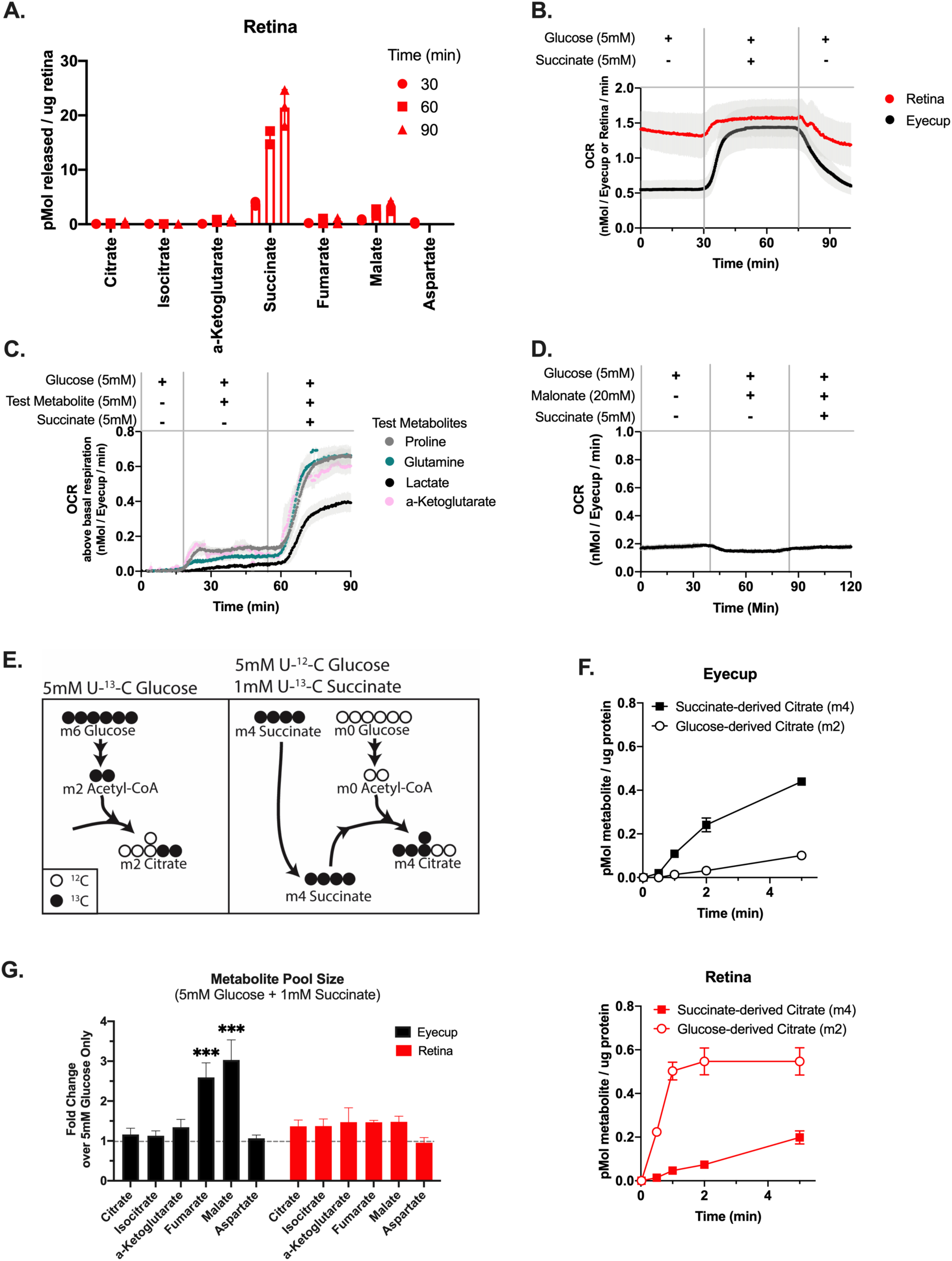
Retinas release succinate which can fuel O_2_ consumption in eyecups. A. TCA cycle metabolites released by retinas incubated in 5 mM ^12^C-glucose. (Error bars indicate S.E.M., n=3 retinas per time point). B. O_2_ consumption of retinas and eyecups perifused with media containing 5 mM glucose, then 5 mM glucose + 5 mM succinate, then 5 mM glucose. Vertical gray bars indicate the approximate time media containing the new metabolite condition reached tissue. (Error bars indicate S.E.M., n=4). C. O_2_ consumption of eyecups supplied with 5 mM glucose, then 5 mM glucose + 5 mM of a test metabolite, then 5 mM glucose + 5 mM test metabolite + 5 mM succinate. Test metabolites are proline (gray), glutamine (teal), lactate (black), and α-ketoglutarate (pink). (Error bars indicate S.E.M., n=3). D. O_2_ consumption of eyecups supplied with 5 mM glucose, then 5 mM glucose + 20 mM malonate, then 5 mM glucose + 20 mM malonate + 5 mM succinate. (Error bars indicate S.E.M., n=2). E. Labeling schematic showing isotopomers of citrate produced by either U-^13^C-glucose alone (left panel) or ^12^C-glucose + U-^13^C-succinate (right panel). F. Citrate production in eyecups (top panel) and retinas (bottom panel) supplied with either 5 mM U-^13^C-glucose alone or 5 mM ^12^C-glucose + 1 mM U-^13^C-succinate. (Error bars indicate S.E.M., n=2 retinas or eyecups per time point). G. Pool size of TCA cycle metabolites in eyecups and retinas supplied with 5 mM ^12^C-glucose + 1 mM U-^13^C-succinate for 10 minutes, relative to tissue supplied with 5 mM^12^C-glucose alone. (Error bars indicate S.E.M., n=2).

The RPE consumes mitochondrial fuels, and RPE cells are in direct contact with the retina and metabolites exchange between these tissues within an eye (Adijanto et al., 2014; Du et al., 2016; Kanow et al., 2017; Nickla and Wallman, 2010; Reyes-Reveles et al., 2017; Swarup et al., 2019; Yam et al., 2019; Young and Bok, 1969). For these reasons, we tested the capacity of RPE cells to consume succinate. We measured O_2_ consumed by intact mouse retinas and eyecups fueled with succinate. For these experiments we used a custom-built perifusion oximeter (Sweet et al., 2002). Succinate causes a 0.6 ± 0.1 nmol/eyecup/min increase in the O_2_ consumption rate (OCR) by eyecups, which is ∼200% greater than OCR by eyecups supplied with 5 mM glucose alone (n=12). Succinate increases OCR in retinas by only 0.2 ±.1 nmol/retina/min, which is ∼15% greater than OCR by retinas supplied with 5 mM glucose alone (n=7) (**Figure 1B**). The K_m_ for eyecup O_2_ consumption when fueled with succinate is 2.1 mM (± 0.1 mM) and the maximal respiration rate is 1.0 nmol/eyecup/min (± 0.4 nmol/eyecup/min) (**Supplemental Figure 1A**). No other glycolytic or TCA cycle metabolite we tested stimulates O_2_ consumption as much as succinate, and the metabolites we tested did not synergistically enhance the succinate-induced OCR (**Figure 1C)**. Addition of the downstream TCA cycle metabolites fumarate and malate partially suppress O_2_ consumption in eyecups stimulated by succinate (consistent with mass action) (**Supplemental Figure 1B**). We confirmed the specificity of this effect to succinate dehydrogenase (SDH), as succinate-stimulated increases in OCR are blocked by the SDH inhibitor malonate (**Figure 1D**).

We used isotopic tracers to determine if the presence of succinate enhances the rate of TCA cycle reactions. We supplied retinas and eyecups with either 5 mM U-^13^C-glucose, or with 5 mM ^12^C-glucose and 1 mM U-^13^C-succinate for times ranging from 0 to 5 minutes. As a readout of TCA cycle flux, we compared the rate of m2 citrate formed from glucose alone (**left panel of Fig. 1E**) with the rate of m4 citrate from succinate (**right panel of Fig. 1E**). Eyecups make citrate from succinate carbons (m4) ∼5-fold faster than from glucose alone (m2) (**Fig. 1F**). In striking contrast, retinas make citrate using carbons from glucose (m2) faster than with carbons from succinate (m4) (**Fig. 1G**). Rates for other TCA cycle reactions are similarly enhanced in eyecups but not retinas (**Supplemental Fig. 1D, E**). Succinate does not alter metabolite pool size in retinas but it is anapleurotic in eyecups, acting to increase fumarate and malate pools in eyecups ∼3-fold relative to when glucose is the only fuel supplied (**Fig. 1H**). Eyecup preparations were used for these experiments because removing the RPE (mechanically or enzymatically) from the eyecups can damage it and alter its metabolic features. Since eyecups contain multiple cell types (RPE, choroidal endothelial, and sclera) we repeated the flux analysis with cultured human fetal RPE (hfRPE) at 2 minutes to assess if RPE uses succinate. These fully differentiated RPE cells make twice as much citrate from succinate carbons (m4) as citrate from glucose carbons (m2) (**Supplemental Fig. 1C**). This suggests that RPE cells are at least one of the types of cells in RPE/choroid preparations that oxidizes succinate.

### Succinate enhances malate export in eyecups

The findings in **Fig. 1** show that retinas can export succinate, and the RPE/choroid in eyecups can use it to fuel mitochondrial respiration. We next investigated the hypothesis that there is a reciprocal exchange of metabolites from the RPE/choroid to the retina. To determine if eyecups release a metabolite that retinas can use to enhance succinate production, we supplied eyecups with 5 mM ^12^C-glucose and quantified TCA cycle metabolites released into the incubation medium at 30, 60 and 90 minutes. Eyecups do not release succinate, but do release α-ketoglutarate, malate, and aspartate (**Figure 2A)**. We then determined if succinate influences the export of any of these metabolites by supplying eyecups with U-^13^C-succinate in addition to ^12^C-glucose (**Figure 2B**). When we provide eyecups with 5 mM ^12^C-glucose and 2 mM U-^13^C-succinate for 30 minutes, they release 15-fold more malate and 271-fold more fumarate compared to when they were supplied with 5 mM glucose alone (**Figure 2C**). The majority of malate and fumarate released is m4, indicating it was made from U-^13^C-succinate. Of the succinate carbons consumed, approximately 43% (± 19%) are released as fumarate and malate.

**FIGURE 2:**
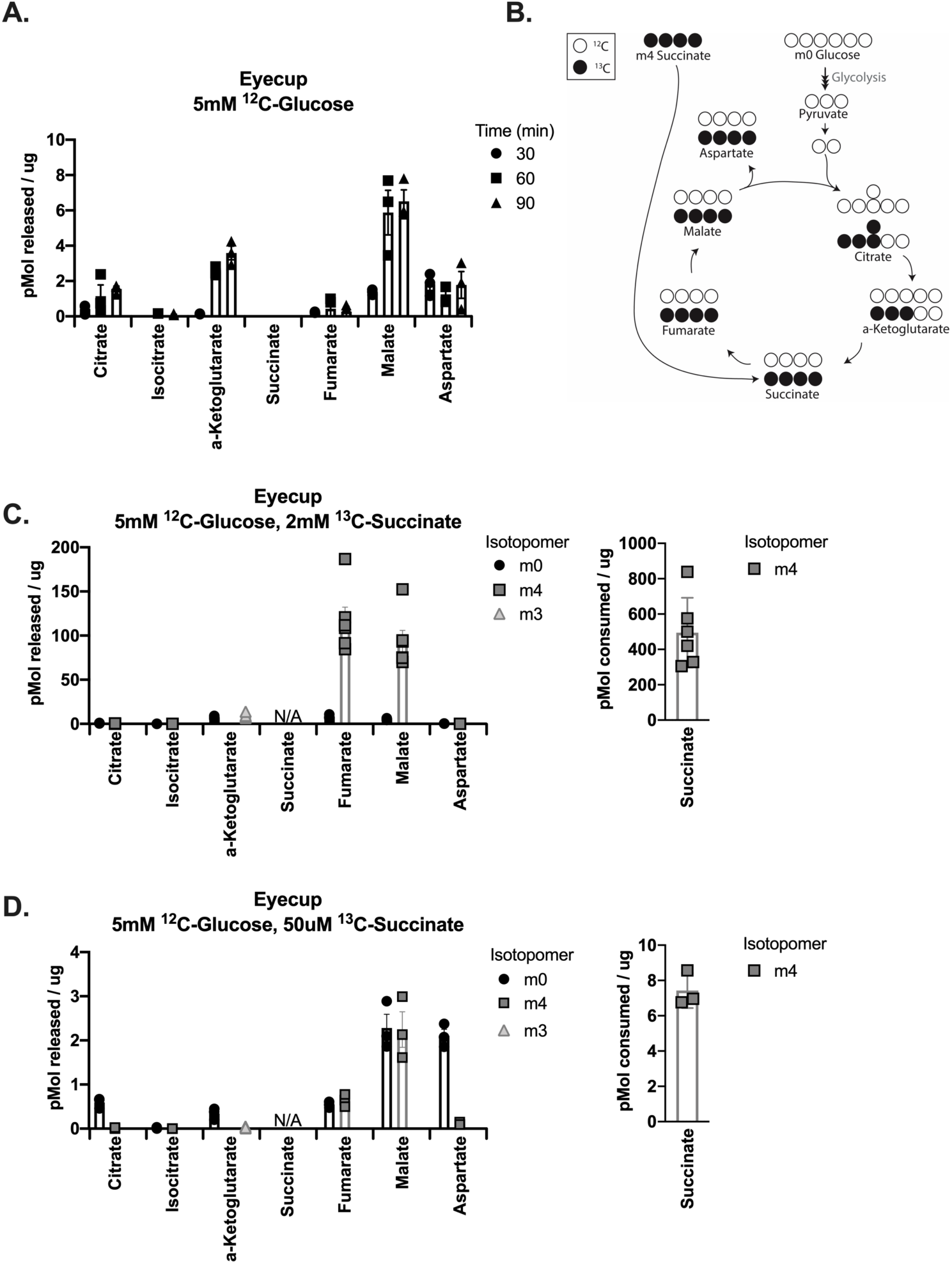
Succinate enhances malate export in eyecups. A. TCA cycle metabolites released by eyecups incubated in 5 mM ^12^C-glucose. (Error bars indicate S.E.M., n=3 eyecups per time point). B. Labeling schematic showing possible TCA cycle isotopomers produced by eyecups supplied with ^12^C-glucose + U-^13^C-succinate. For simplicity, only isotopomers produced in a single turn of the TCA cycle are shown. C. TCA cycle metabolites released by eyecups incubated in 5 mM ^12^C-glucose + 2 mM U-^13^C-succinate for 30 minutes. Succinate consumed by eyecups during incubation is shown on the right panel. (Error bars indicate S.E.M., n=6 eyecups). D. TCA cycle metabolites released by eyecups incubated in 5 mM ^12^C-glucose + 50 μM U-^13^C-succinate for 30 minutes. Succinate consumed by eyecups during incubation is shown on the right panel. (Error bars indicate S.E.M., n=3

We cannot determine the concentration of succinate that RPE cells are exposed to from a retina in the context of a living eye, so we also performed this experiment with a much lower concentration of succinate. When eyecups are supplied with 5 mM ^12^C-glucose and 50 μM U-^13^C-succinate for 30 minutes, they export 3-fold more malate and 6-fold more fumarate compared to when they were supplied with 5 mM glucose alone (**Figure 2D**). Similarly to high succinate exposure, 39% (± 11%) of the succinate carbons are released as fumarate and malate. Remarkably, eyecups export equivalent amounts of m4 malate (made from succinate) and m0 malate (made from glucose). This is despite being incubated in a 100-fold lower concentration of succinate compared to glucose. Incubating retinas in 5 mM ^12^C-glucose and 50 μM U-^13^C-succinate for 30 minutes does not similarly enhance fumarate and malate export, and retinas do not measurably deplete succinate from the incubation media (**Supplemental Figure 2A, 2B).**

### Retinas can use malate to produce succinate via reverse electron transport at SDH

Succinate enhances the export of malate and fumarate from the RPE/choroid complex (**Fig 2**). Even at low concentrations of succinate, eyecups export significant quantities of malate. We next tested whether malate exported by eyecups can influence retinal metabolism. Retinas in an eye are in a hypoxic environment, and hypoxic tissues can produce succinate by both canonical oxidative TCA cycle activity as well as through reverse electron transport at SDH (Chouchani et al., 2014; Zhang et al., 2018). Since retinas export succinate (**Fig. 1**), they require an anaplerotic source of metabolites feeding into the TCA cycle to avoid running out of TCA cycle intermediates. Based on this premise, we tested if retinas might use malate for anaplerosis to sustain succinate synthesis by reverse electron transport at SDH.

We used U-^13^C-malate to quantify reverse electron transport at SDH in both retinas and eyecups (**Figure 3A**). We incubated retinas and eyecups for 5 minutes in U-^13^C-malate at a range of concentrations (5, 50, and 500 μM), all in the presence of 5 mM ^12^C-glucose. At all concentrations, retinas form much more m4 succinate than eyecups (**Figure 3B, 3C**). None of these concentrations of malate increase the size of the fumarate pool in retinas (**Figure 3C**). This shows that the formation of m4 succinate we observe in retinas is not driven only by an increase in concentration of the reactant of the reverse SDH reaction (fumarate) and is instead a property that is inherent to retinas and not eyecups. Total metabolite levels for other metabolites are reported in **Supplemental 3A.** The formation of m4 succinate by retinas supplied with 50 μM U-^13^C-malate is blocked by 20 mM of the SDH inhibitor malonate (**Figure 3D**, other m4 metabolites shown **Supplemental Figure 3B**).

**FIGURE 3:**
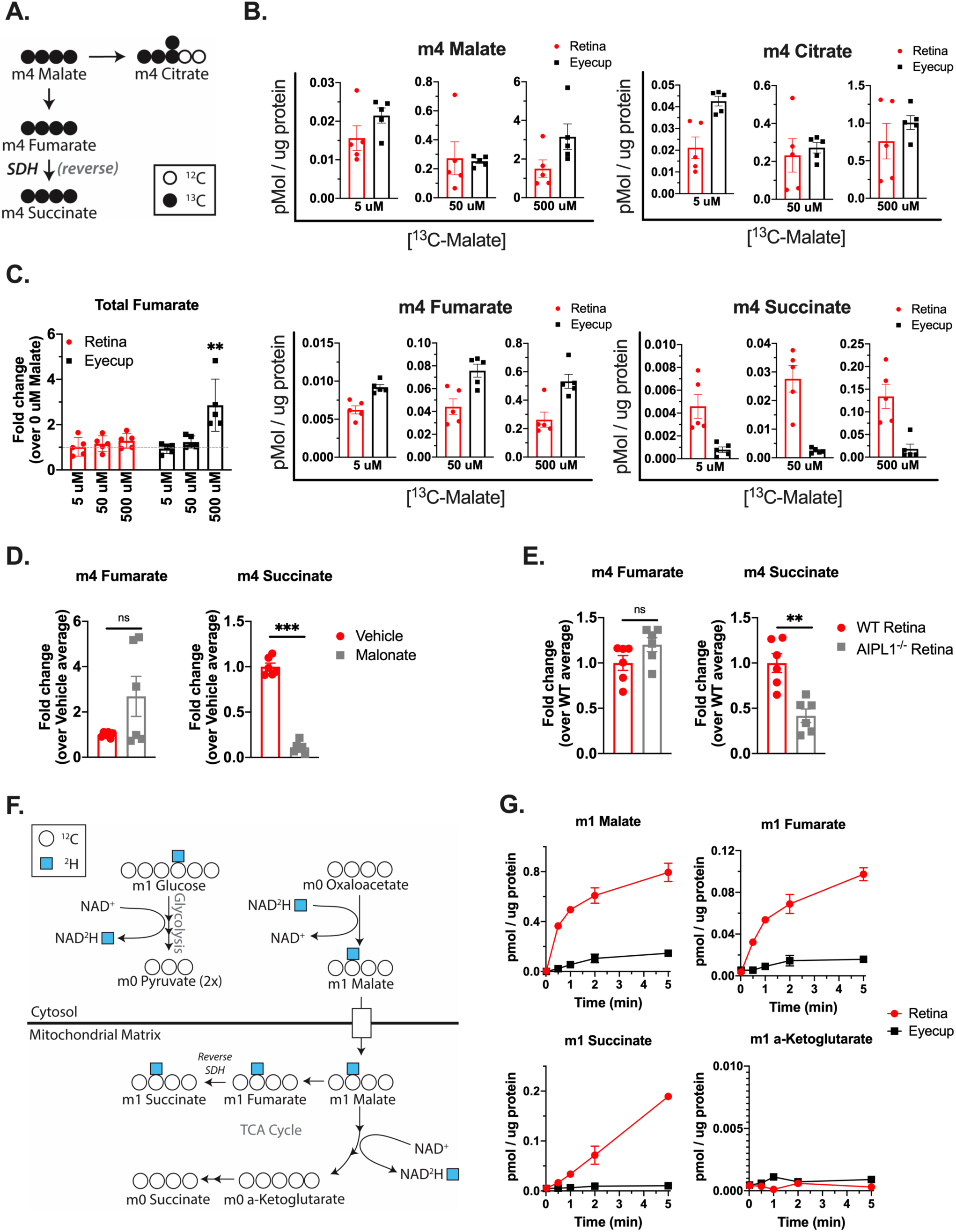
Retinas can use malate to produce succinate via reverse electron transport at SDH. A. Labeling schematic showing isotopomers of TCA cycle metabolites produced by tissue incubated in U-^13^C-malate + ^12^C-glucose. B. Isotopomers produced by retinas and eyecups incubated in 5 mM ^12^C-glucose + 5, 50, or 500 μM U-^13^C-malate for 5 minutes. (Error bars indicate S.E.M., n=5 retinas or eyecups per concentration). C. Total fumarate levels in retinas and eyecups incubated in 5 mM ^12^C-glucose + 5, 50, or 500 μM U-^13^C-malate for 5 minutes, relative to tissue incubated in 5 mM ^12^C-glucose alone (gray line). (Error bars indicate S.E.M., n=5 retinas or eyecups per concentration). D. m4 fumarate and m4 succinate produced by retinas supplied with 5 mM ^12^C-glucose + 50 μM U-^13^C-malate + 20 mM Malonate for 5 minutes. (Error bars indicate S.E.M., n=3 retinas). E. m4 fumarate and m4 succinate produced by WT and AIPL1^-/-^ retinas supplied with 5 mM ^12^C-glucose + 50 μM U-^13^C-malate for 5 minutes. (Error bars indicate S.E.M., n=6 retinas). F. Labeling schematic showing isotopomers produced by 4-^2^H-glucose. G. Accumulation of deuterated (m1) malate, fumarate, succinate α-ketoglutarate in retinas and eyecups incubated in 5 mM 4-^2^H-glucose for 0.02, 0.5, 1, 2, and 5 minutes. (Error bars indicate S.E.M., n=2 retinas per time point).

Retinas are composed of several types of neurons, so we next investigated if a specific cell type in the retina favors reverse SDH activity. Photoreceptors are the neurons in the lowest pO_2_ layer of the mouse retina and their outer segments are in direct contact with the RPE in an intact eye, so they are a candidate site of reverse SDH activity. To determine if photoreceptors favor reverse SDH activity, we supplied AIPL1^-/-^ retinas (in which photoreceptors have degenerated and are no longer present) with 5 mM ^12^C-glucose and 50 μM U-^13^C-malate for 5 minutes (Dyer et al., 2004). AIPL1^-/-^ retinas produce equivalent amounts of m4 fumarate but less than half as much m4 succinate as WT retinas per μg of protein (**Figure 3E**, other m4 metabolites shown **Supplemental Figure 3C**). This indicates that the contribution of photoreceptors to reverse SDH activity is greater than that of the inner retina. We also tested if photoreceptors specifically release succinate by incubating WT and AIPL1^-/-^ retinas in 5 mM ^12^C-glucose and 50 μM U-^12^C-malate for 30 minutes and measuring exported TCA cycle metabolites (**Supplemental Figure 3D**). We found that retinas lacking photoreceptors release less succinate and more fumarate per μg of protein, confirming that photoreceptors are a major contributor to succinate release.

When supplying U-^13^C-malate to eyecups, we observed that we could “force” SDH to operate in reverse if we supplied enough U-^13^C-malate (500 μM) to cause the total amount of fumarate to increase (**Figure 3C**). For this reason, we sought to also measure reverse SDH activity in a manner that does not perturb the size of the fumarate pool, i.e. in the presence of glucose alone. We used 4-^2^H-glucose to track reverse electron transport in the absence of added mitochondrial fuels. When cells metabolize 4-^2^H-glucose, the deuterium is transferred from the carbon skeleton to NADH during the GAPDH reaction in glycolysis (**Figure 3F)** (Lewis et al., 2014). This NAD^2^H then can be used in cytosolic reactions, including reduction of oxaloacetate to malate. Deuterated malate then can enter mitochondria. Since the fumarase reaction is readily reversible, deuterated malate will equilibrate with the fumarate pool. In the case of reverse SDH activity, deuterated fumarate will be converted to deuterated succinate. We incubated retinas and eyecups in 5 mM 4-^2^H-glucose for times ranging from 0 to 5 min and observed steady accumulation of m1 (deuterated) succinate in retinas but not in eyecups **(Figure 3G)**. This further indicates that retinas but not eyecups exhibit reverse SDH activity. To ensure that the deuterated succinate we observe is produced solely from reverse SDH activity, we confirmed that the deuterium was transferred off of the carbon skeleton during oxidative TCA cycle activity prior to the formation of α-ketoglutarate (**Figure 3G**).

### Reverse SDH activity maintains a significant portion of the succinate pool in retinas

It has been estimated that reverse SDH activity produces only about ∼6% of succinate during ischemia in heart (Zhang et al., 2018). To determine if reverse SDH activity produces a more substantial amount of succinate in retina, we next quantified the contributions of reverse SDH activity and canonical TCA cycle activity to maintaining the retinal succinate pool.

To compare most directly the different modes of succinate production under the same conditions, we used U-^13^C-glucose to track oxidative TCA cycle activity and 4-^2^H-glucose to track reverse SDH activity. To assess the contribution of oxidative TCA cycle activity to succinate production, we incubated retinas in 5 mM U-^13^C-glucose for times ranging from 0 to 60 minutes and determined the fractional enrichment of m2 succinate at the steady-state by fitting a curve assuming a first order reaction (**Figure 4A**). At the steady-state, 6.9% of the succinate pool is m2 in retinas. However, this does not mean that only 6.9% of the succinate pool is maintained by oxidative TCA cycle activity, since its oxidative precursor α-ketoglutarate is only 22.8% m2-labeled at the steady-state. In order to account for incomplete labeling of the α-ketoglutarate pool, we scaled these values to determine what the fractional enrichment of m2 succinate would be if 100% of the α-ketoglutarate pool were m2 (i.e. 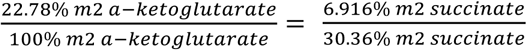). This calculation predicts that the fraction of the succinate pool that is formed from α-ketoglutarate (via oxidative TCA cycle activity) is 30.4%.

**FIGURE 4:**
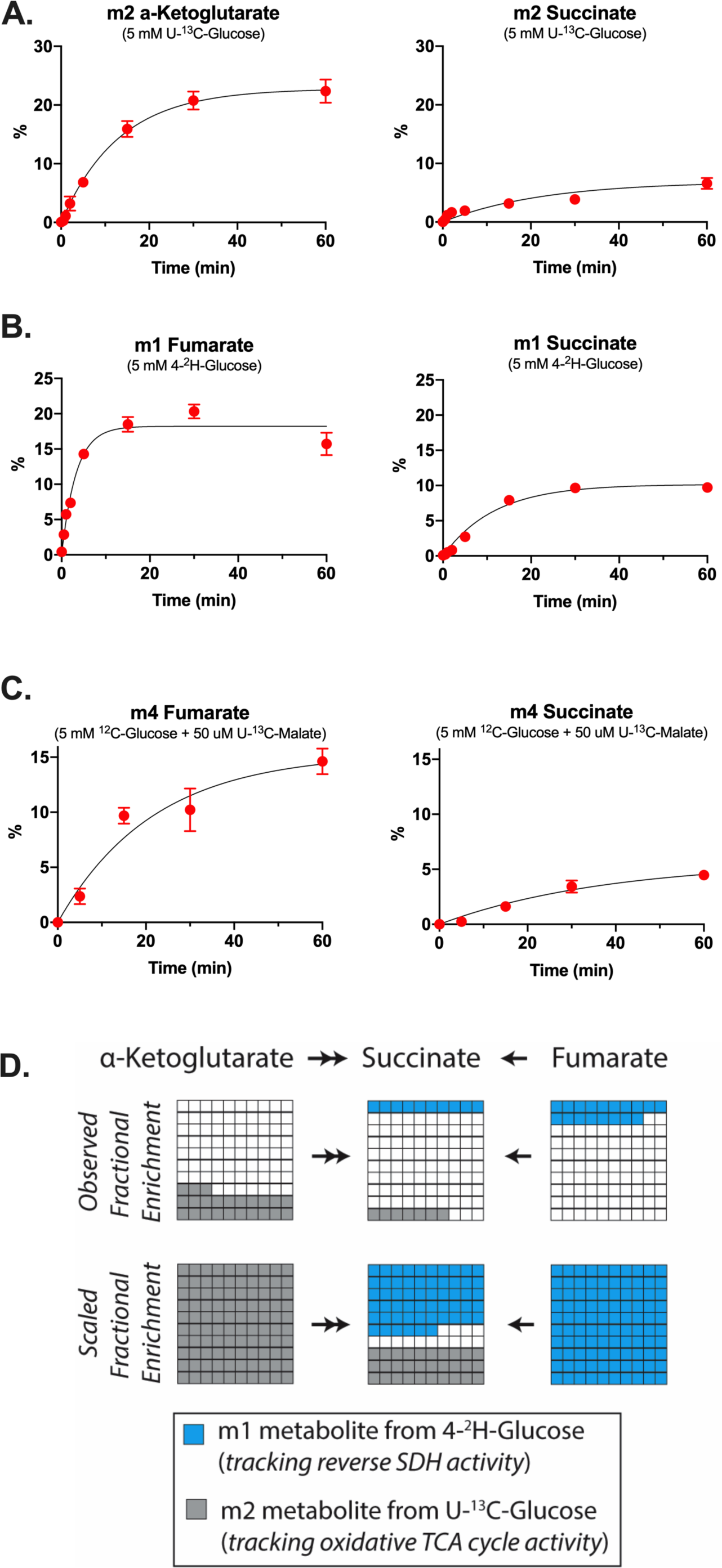
Reverse SDH activity maintains a significant portion of the succinate pool in retinas. A. Fractional enrichment of m2 α-ketoglutarate and m2 succinate in retinas supplied with 5 mM U-^13^C-glucose for 0.02, 0.5, 1, 2, 5, 15, 30, and 60 minutes. (Error bars indicate S.E.M., n=4 to 7 retinas per time point). B. Fractional enrichment of m1 fumarate and m1 succinate in retinas supplied with 5 mM 4-^2^H-glucose for 0.02, 0.5, 1, 2, 5, 15, 30, and 60 minutes. (Error bars indicate S.E.M., n=3 retinas per time point). C. Fractional enrichment of m4 fumarate and m4 succinate in retinas supplied with 5 mM ^12^C-glucose + 50 μM U-^13^C-malate for 0.02, 5, 15, 30, and 60 minutes. (Error bars indicate S.E.M., n=3 to 5 retinas per time point). D. Schematic showing relative contributions of oxidative TCA cycle activity and reverse SDH activity to succinate production in retinas supplied with glucose only. Observed fractional enrichments are shown in the top panel, and fractional enrichments scaled for complete labeling of reactant pools are shown in the bottom panel.

To measure directly the contribution of reverse SDH activity to succinate production when glucose is the only fuel source available, we repeated this experiment under the same conditions using 4-^2^H-glucose. At the steady-state, the fractional enrichment of m1 fumarate is 18.2% and m1 succinate is 10.2% (**Figure 4B**). Scaling these values for 100% m1 labeling of the fumarate pool predicts that reverse SDH activity maintains 55.8% of the succinate pool. m1 α-ketoglutarate is not detected at any time, indicating that m1 succinate does not form by oxidative TCA cycle activity (**Supplemental Figure 4A**).

We also determined what fraction of the retinal succinate pool is maintained by reverse SDH activity from exogenous malate. We incubated retinas in 50 μM U-^13^C-malate and 5 mM ^12^C glucose for times ranging from 0 to 60 minutes (**Figure 4C**). The steady-state fractional enrichment was calculated as 15.3% for m4 fumarate and 5.7% for m4 succinate. Scaling these values for 100% m4 labeling of the fumarate pool predicts that 37.2% of the succinate pool is maintained from exogenous malate. This value is lower than the fraction of the succinate pool maintained by reverse SDH activity when glucose is the only fuel. Since the retina contains many cell types, a possible explanation for this difference is that there is a population of cells in the retina that consumes malate (producing m4 fumarate) but does not produce m4 succinate.

In summary, our analysis in retinas supplied with glucose alone (i.e. under identical conditions) predicts that 30.4% of retinal succinate is maintained by oxidative TCA cycle activity and 55.8% is maintained by reverse SDH activity (**Figure 4D**). All steady-state fractional enrichment values and reaction constants with confidence intervals are reported in **Supplemental Figure 4B**. Fractional enrichment of all isotopomers for relevant metabolites at 60 minutes are reported in **Supplemental Figure 4C, D**, and **E**.

### Low COXIV expression drives reversal of SDH in retinas

**Figures 3** and **4** show that reverse electron transport at SDH is a predominant pathway for succinate generation in retinas and not in eyecups. This led us to investigate possible molecular differences between these two tissues that could drive this specialization. Reduced ubiquinone (QH_2_) in the mitochondrial inner membrane normally donates electrons to reduce O_2_ to H_2_O via complex IV. However, we observe in retinas that electrons from QH_2_ are used to reduce fumarate to succinate, even in retinal explants assayed in air with 21% O_2_. Since this indicates that lack of O_2_ alone does not drive reversal of SDH, we hypothesized that mitochondria in retinas might have limiting complex IV, thus leading to an accumulation of QH_2_ which then drives reversal of SDH even during normoxia.

We evaluated expression of ETC component protein levels in retinal and eyecup homogenates by immunoblotting with a mix of antibodies that recognize representative subunits of the ETC complexes (**Fig 5A**, lanes labeled “Fresh”). We found that freshly dissected retinas have a lower ratio of COXIV (a component of complex IV) to ATP5A (a component of ATP synthase) compared to eyecups. Hypoxia has been observed to decrease COXIV protein levels (Fukuda et al., 2007; Vijayasarathy et al., 2003). This motivated us to evaluate the influence of O_2_ levels on COXIV expression in retinas and eyecups. We incubated retinas at 0%, 1%, 5%, 21% and 95% pO_2_ for 2 hours in 5 mM glucose and then analyzed ETC component expression by immunoblotting. The low COXIV/ATP5a ratio in freshly dissected retinas is similar to the ratio in retinas cultured in 0% O_2_. Increasing pO_2_ leads to a higher COXIV/ATP5A ratio in retinas. **Figure 5A** shows a representative immunoblot, and quantification of multiple independent experiments is shown in panel **B**. Altering pO_2_ does not have a substantial effect on the COXIV/ATP5A ratio in eyecups. These results indicate that the hypoxic niche retinas reside in suppresses COXIV expression and thus complex IV activity.

**FIGURE 5:**
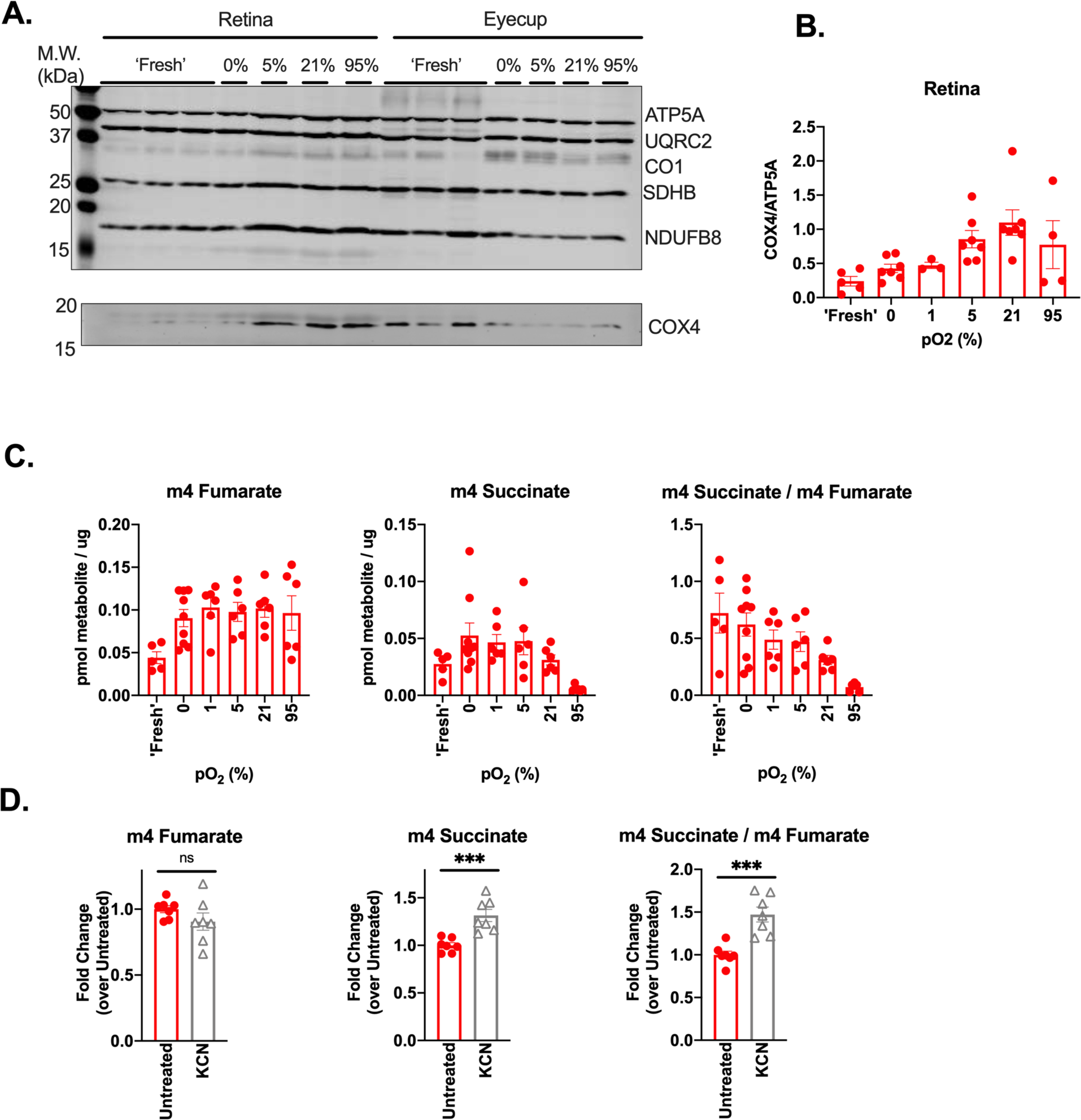
Low COXIV expression drives reversal of SDH in retinas. A. Immunoblot of retinas and eyecups using an ETC component cocktail (top blot) and a single antibody against COX4 (bottom blot). ATP5A: ATP synthase subunit 5A, UQRC2: ubiquinol cytochrome c oxidoreductase subunit core 2, CO1: cytochrome *c* oxidase subunit 1, SDHB: succinate dehydrogenase B, NDUFB8: NADH Ubiquinone Oxidoreductase Subunit B8, COX4: cytochrome *c* oxidase subunit 4. B. Quantification of multiple immunoblots probed with COX4 and ATP5A. (Error bars indicate S.E.M., n=3-7 retinas at each O_2_ concentration). C. m4 fumarate, m4 succinate, and the m4 succinate/m4 fumarate ratio measured in retinas supplied with 5 mM ^12^C-glucose + 50 μM U-^13^C-malate for 5 minutes after pre-equilibration at specified pO_2_ for 2h. ‘Fresh’ data is duplicated from experiments in figure 3b for ease of comparison (Error bars indicate S.E.M., n=6-9). D. m4 fumarate, m4 succinate, and the m4 succinate/m4 fumarate ratio measured in retinas supplied with 5 mM ^12^C-glucose + 50 μM U-^13^C-malate in the presence of KCN. (Error bars indicate S.E.M., n=6).

Based on these observations, we hypothesized that raising COXIV expression by pre-incubating retinas with higher O_2_ levels would diminish the conversion of fumarate to succinate. To test this, we preconditioned retinas and eyecups for 2 hours in pre-equilibrated media containing 5 mM ^12^C-glucose at varied pO_2_. After 2 hours, reverse SDH activity was assessed by transferring retinas into fresh media containing 5 mM ^12^C-glucose and 50 μM U-^13^C-malate and incubating at the same pO_2_ as before for 5 minutes. Retinas incubated in increasing pO_2_ exhibit less reverse SDH activity, as measured by both m4 succinate production and by the m4 succinate/m4 fumarate ratio (**Figure 5C**) However, we also noted that pre-equilibration for 2h (at any pO_2_) drastically alters the total succinate pool size in retinas (**Supplemental Figure 5A**).

Pre-incubating retinas at increasing pO_2_ increases COXIV levels and also decreases reverse SDH activity. However, changes in total metabolite levels suggest that pre-incubation also causes other metabolic changes which are independent of COXIV but could also influence reversal of SDH. For this reason, we also used pharmacological means to more directly interrogate the influence of complex IV on reverse SDH activity. We hypothesized that treating freshly dissected retinas with the complex IV inhibitor KCN would further increase the reduction state of the retinal Q pool and drive more reverse SDH activity. We incubated retinas in 5 mM ^12^C-glucose and 50 μM U-^12^C-malate with 3 mM KCN for 5 minutes, then transferred the retinas to fresh media containing 5 mM ^12^C-glucose and 50 μM U-^13^C-Malate with 3 mM KCN for 5 minutes to assay reverse SDH activity. We found that retinas treated with KCN produce more m4 succinate, which is consistent with a buildup of QH_2_ further driving reverse SDH activity (**Figure 5D**). We also observed that KCN treatment depleted the total fumarate pool in retinas (**Supplemental Figure 5B**). Together, these results show that modulating complex IV activity can directly influence reverse SDH activity in retinas.

## DISCUSSION

We have found evidence for a succinate/malate cycle between the mitochondria of the retina and RPE/choroid. In this cycle, the retina releases succinate, the RPE/choroid imports the succinate to fuel mitochondrial respiration and in turn exports malate, which refuels the retina. Low COXIV expression appears to bolster reverse SDH activity in retinas, likely as a consequence of increasing the reduction state of ubiquinone (Q) (**Fig. 6A, B**). We find that pO_2_ influences expression of COXIV in retinal explants, suggesting that it is the hypoxic environment of the retina that keeps complex IV activity low.

**Fig. 6.**
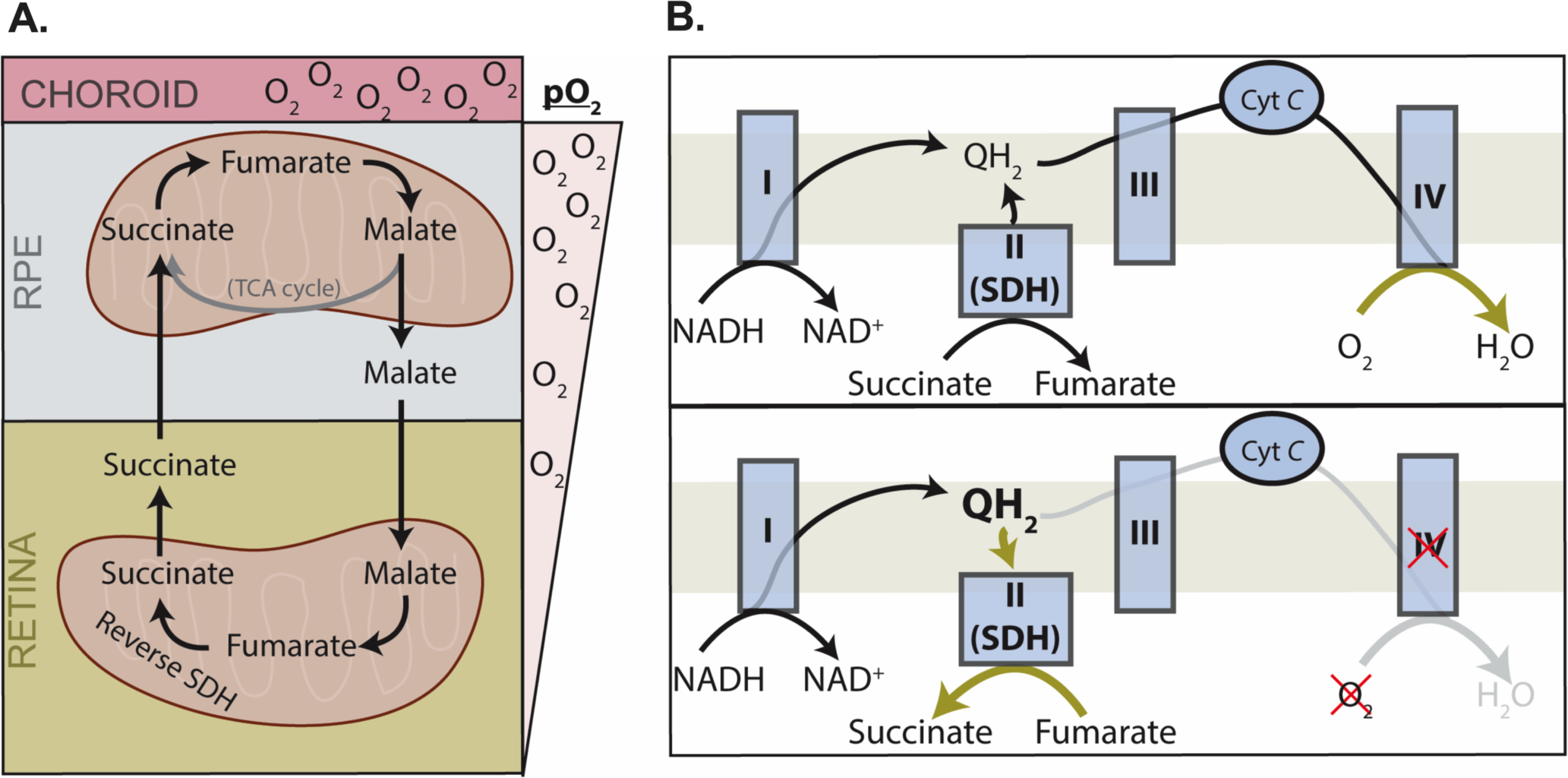
Schematic representation of succinate and malate exchange in the retina and RPE. A. The malate/succinate shuttle that transfers reducing power from the hypoxic retina to the O_2_-rich RPE/choroid complex. B. Normal electron transport chain function (upper panel) vs. retinal electron chain function (lower panel). Low COXIV expression in the retina causes SDH to act as a release valve, transferring electrons from QH_2_ onto fumarate to produce succinate.

Reverse electron transport at SDH is a major pathway for succinate production in the retina. Previous studies have shown that reverse electron transport can produce succinate in the ischemic heart; however, it was calculated that only 5.7% of the succinate pool was made by reverse SDH activity (Chouchani et al., 2014; Zhang et al., 2018). In contrast, we observe that as much as 55% of the succinate pool in retinas derives from reverse SDH activity. Remarkably, this is greater than the amount of succinate produced from oxidative TCA cycle activity.

The retina relies heavily on glycolysis to produce ATP (Kanow et al., 2017; Krebs, 1927). Since the RPE lies between the retina and the choroidal blood supply, glucose must pass through the RPE mostly unconsumed in order to fuel glycolysis in the retina. Export of succinate from the retina to the RPE provides the RPE with an alternative fuel source to glucose. This allows a greater fraction of glucose to pass through the RPE unconsumed so that it can reach the retina. This suggests that the RPE has specifically adapted to consume succinate, since most tissues (except brown fat) are thought to be impermeable to succinate (Ehinger et al., 2016; Hems et al., 1968; Mills et al., 2018).

Succinate exported from the retina acts as an electron shuttle, carrying reducing power to the O_2_-rich RPE where the electrons can be better used to generate cellular energy. This is not the first succinate-mediated ‘redox shuttle’ to be proposed. During whole-body hypoxia in rats, succinate released from peripheral tissues has also been hypothesized to carry unused reducing power to the lungs, where O_2_ is relatively more accessible during hypoxia (Cascarano et al., 1976).

RPE consumption of succinate can protect the retina by preventing unwanted succinate accumulation. In an oxygen-induced retinopathy model, rats were exposed to 24-hour cycles of hypoxia and hyperoxia from P0 to P21, which caused a 3-fold increase in retinal succinate. Accumulated succinate signaled through GPR91 in retinal ganglion cells to induce pathological extraretinal neovascularization (Sapieha et al., 2008). We have shown that retinas constitutively release succinate. If the RPE were not also constitutively consuming this succinate, it could accumulate in the retina and stimulate unwanted angiogenesis.

The 3-fold stimulation of O_2_ consumption by succinate in the RPE/choroid could also reflect a mechanism that protects the retina from oxidative damage. Mammalian retinas are composed of terminally differentiated neurons that cannot be replaced when damaged. Reactive oxygen species pose a great risk to these neurons since they can damage proteins, membranes, and nucleic acids. However, as long as photoreceptors release succinate, it can stimulate the RPE to consume a significant portion of O_2_ from the choroidal blood supply, thus preventing O_2_ from forming ROS in the retina. This is supported by the observation that although mouse eyecups contain approximately 3-fold less cellular material than retinas, they are able to consume as much O_2_ as the entire retina when stimulated by succinate (**Figure 1B**).

The ecosystem formed by succinate-malate exchange between the retina and RPE illustrates another way that photoreceptor degeneration can drastically impact overall eye health. In the absence of photoreceptors, we observe that retinal succinate export decreases (**Supplemental Figure 3D**). We expect that in an intact eye, this will lead to a decrease in RPE O_2_ consumption. If succinate-stimulated RPE O_2_ consumption normally protects the retina, a loss of photoreceptors would exacerbate oxidative damage to the retina. This may occur in retinitis pigmentosa, where degeneration of rod photoreceptors causes an increase in photoreceptor layer O_2_ tension and leads to secondary cone photoreceptors death (Campochiaro and Mir, 2018; Yu et al., 2000). It may be possible to prevent secondary cone degeneration by supplying exogenous succinate to the RPE, which could recapitulate a critical aspect of this metabolic ecosystem for therapeutic benefit.

Overall, our findings suggest that the retina has adapted to its hypoxic niche by altering the stoichiometry of its respiratory complexes to favor reverse SDH activity. This adaptation can divert electrons out of the electron transport chain to reduce fumarate to succinate. The succinate released from the retina shuttles electrons to the RPE/choroid where they reduce O_2_ to H_2_O. This exchange of metabolites facilitates transport of electrons away from a tissue that is not well poised to transfer them to O_2_ them (the retina) to one that is better suited to use them to reduce O_2_ to H_2_O (the RPE/choroid) (**Fig. 6A, B**). Succinate stimulates RPE cells to release malate as “empty” electron shuttles, which retinas then refill with electrons via reverse SDH activity. This succinate/malate exchange is another aspect of the metabolic ecosystem formed by the retina, RPE, and choroid; where tissue specialization enhances survival in a hazardous and competitive metabolic environment.

## SUPPLEMENTAL FIGURES

**SUPPLEMENTAL FIGURE 1.**
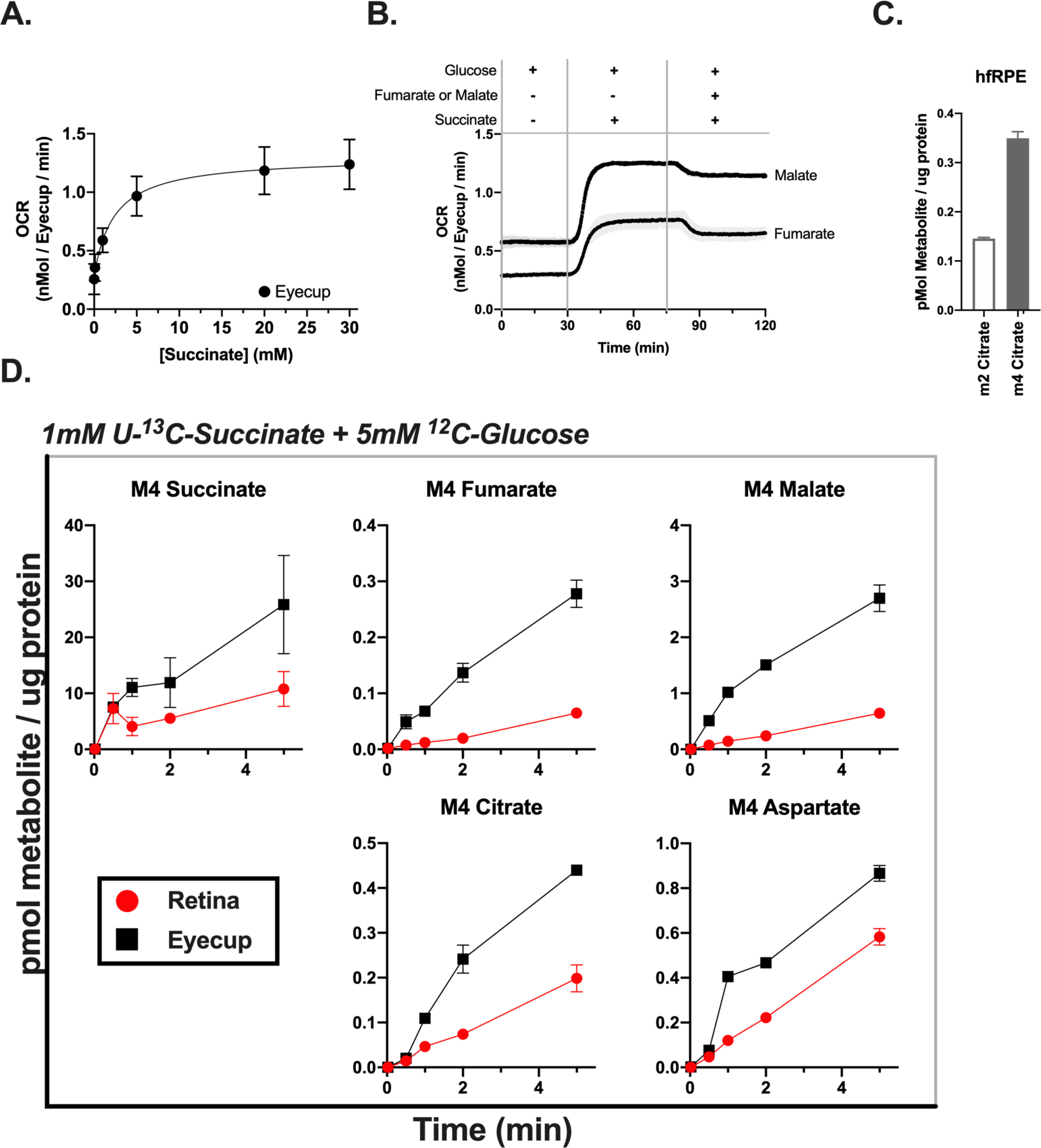
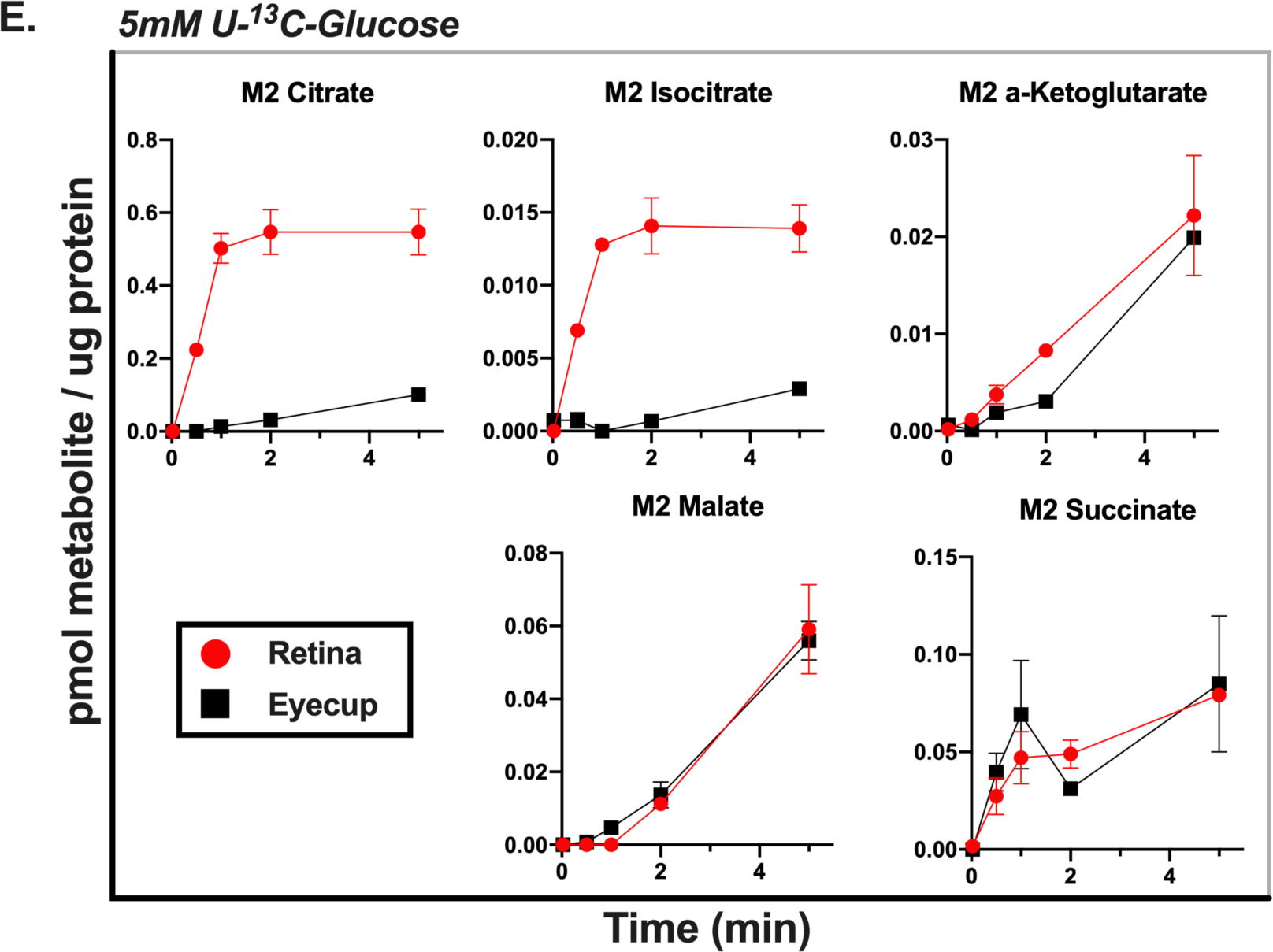
A. Oxygen consumption of eyecups perifused with varying amounts of succinate (0, 0.1, 1, 5, 20, and 30 mM) in the presence of 5 mM glucose. (Error bars indicate S.E.M., n=3). B. Oxygen consumption of eyecups perifused with media containing 5 mM glucose, then 5 mM glucose + 5 mM fumarate or malate, then 5 mM glucose + 5 mM fumarate or malate + 5 mM Succinate. (Error bars indicate S.E.M., n=2 chambers). C. Citrate production in human fetal RPE (hfRPE) supplied with either 5 mM U-^13^C-glucose alone (m2 Citrate) or 5 mM ^12^C-glucose + 1 mM U-^13^C-succinate (m4 Citrate) for 2 minutes. (Error bars indicate S.E.M., n=2). D. Accumulation of TCA cycle metabolites derived from succinate in retinas and eyecups supplied with 5 mM ^12^C-glucose + 1 mM U-^13^C-succinate. (Error bars indicate S.E.M., n=2 retinas or eyecups per time point). E. Accumulation of TCA cycle metabolites derived from glucose in retinas and eyecups supplied with 5 mM ^13^C-glucose. (Error bars indicate S.E.M., n=2 retinas or eyecups per time point).

**SUPPLEMENTAL FIGURE 2.**
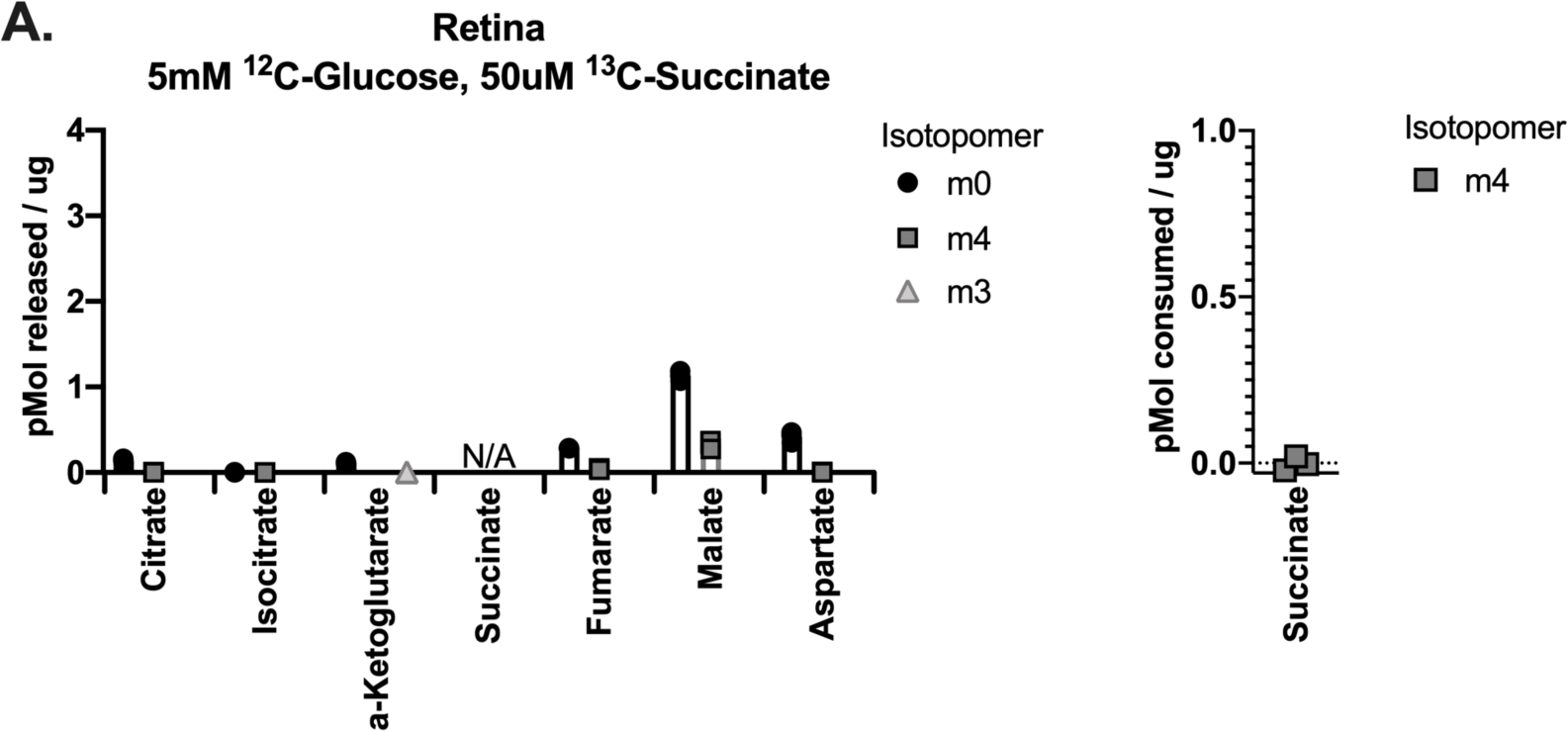
A. TCA cycle metabolites released by retinas incubated in 5 mM ^12^C-glucose + 50 μM U-^13^C-succinate for 30 minutes. Succinate consumed by retinas during incubation is shown on the right panel. (Error bars indicate S.E.M., n=3 retinas).

**SUPPLEMENTAL FIGURE 3.**
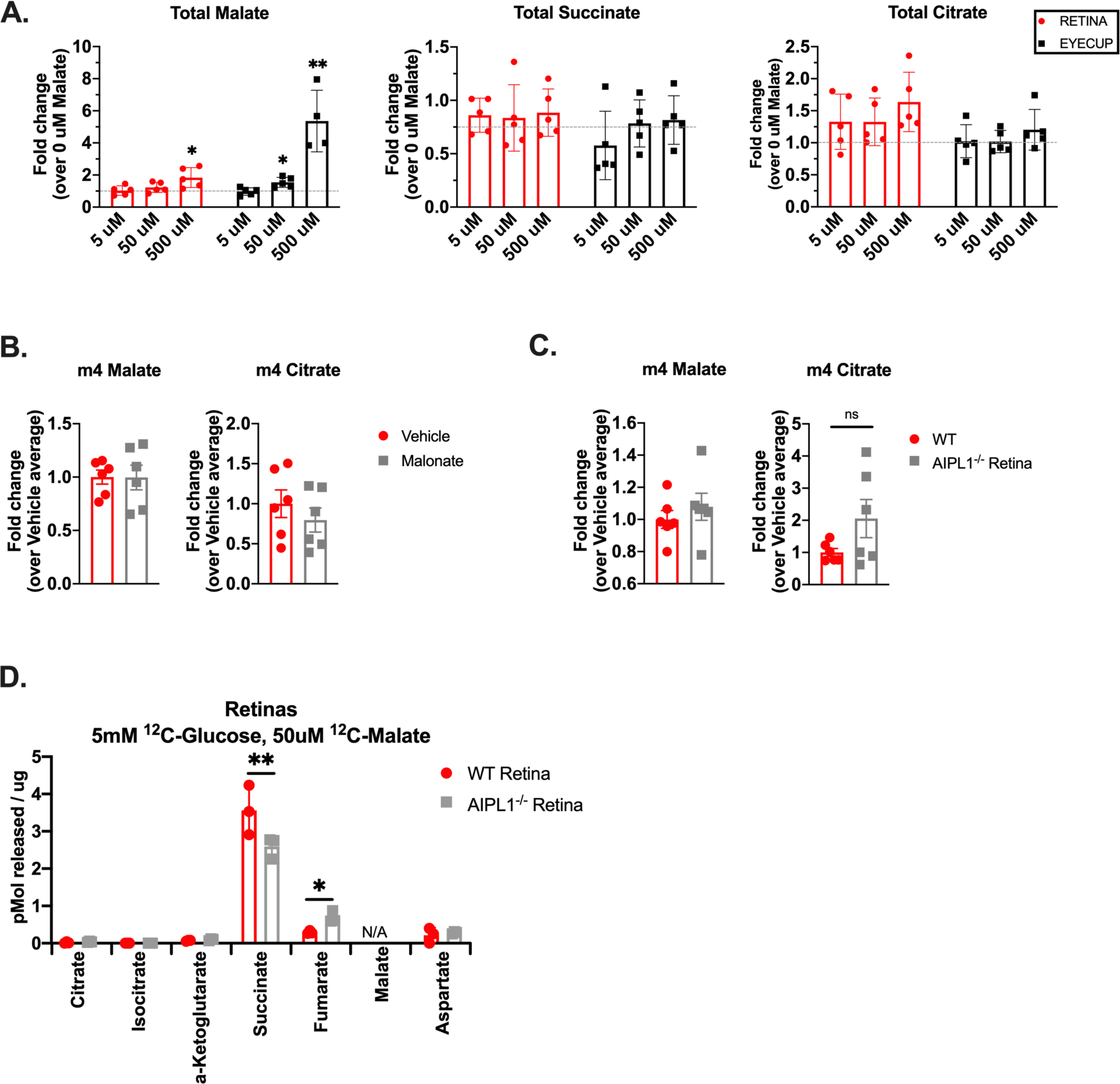
A. Total malate, succinate, and citrate levels in retinas and eyecups incubated in 5 mM^12^C-glucose + 5, 50, or 500 μM U-^13^C-malate for 5 minutes. (Error bars indicate S.E.M., n=5 retinas or eyecups per concentration). B. m4 malate and m4 citrate levels in retinas incubated in malonate (relative to vehicle). (Error bars indicate S.E.M., n=6 retinas). C. m4 malate and m4 citrate levels in AIPL1^-/-^ retinas (relative to control). (Error bars indicate S.E.M., n=6 retinas). D. TCA cycle metabolites released by WT and AIPL1^-/-^ retinas incubated in 5 mM ^12^C-glucose + 50 μM U-^13^C-Malate for 30 minutes. (Error bars indicate S.E.M., n=3 retinas).

**SUPPLEMENTAL FIGURE 4.**
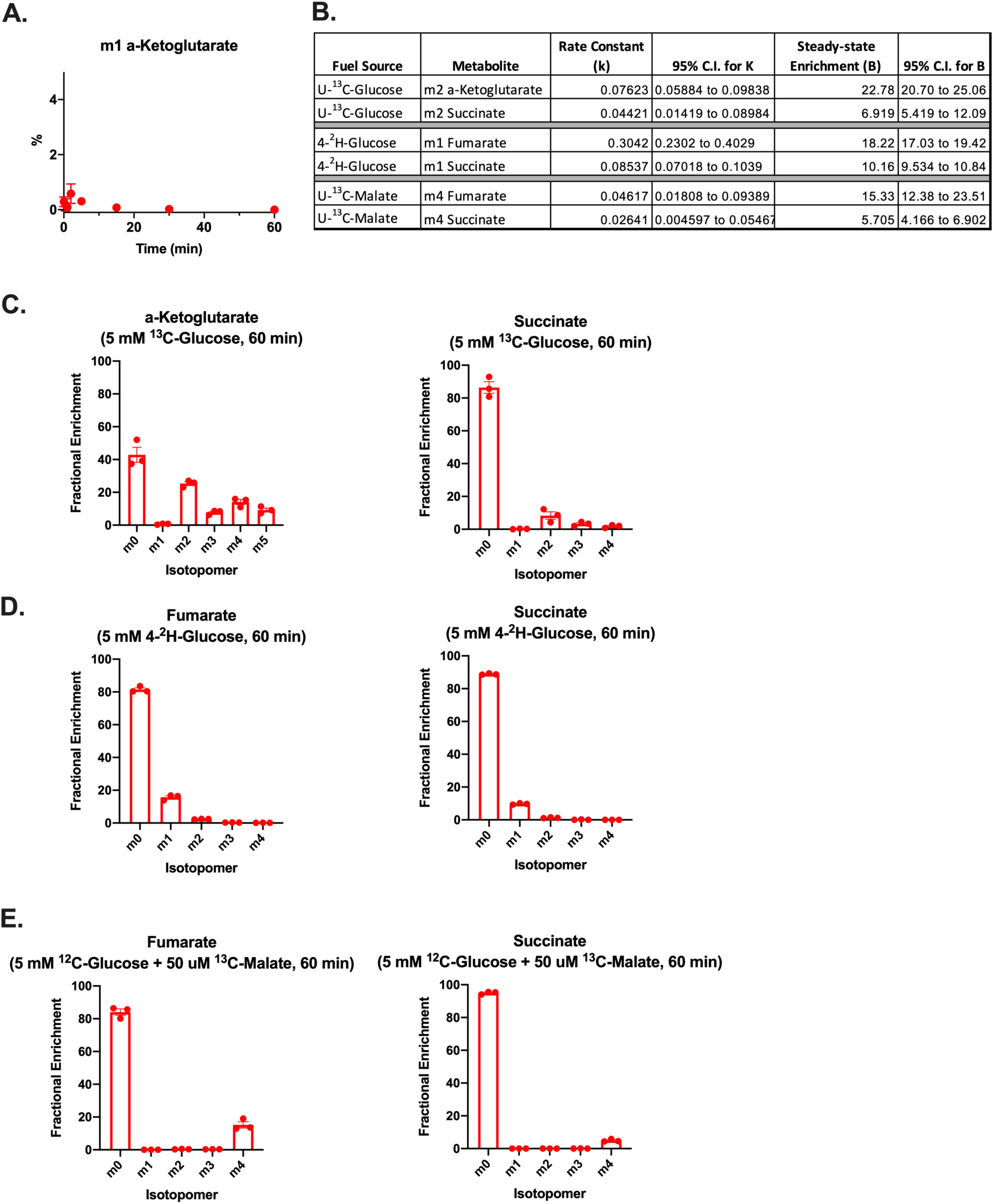
A. Fractional enrichment of m1 a-ketoglutarate in retinas supplied with 5 mM 4-^2^H-glucose for 0.02, 0.5, 1, 2, 5, 15, 30, and 60 minutes. (Error bars indicate S.E.M., n=3 retinas per time point). B. Table of constants and confidence intervals calculated by Prism using the equation % *metabolite* = *B* ∗ (1 – *e*^−*kt*^) to fit each curve. B is the fractional enrichment at the steady state, k is the rate constant for the reaction, t is time, and %metabolite is the fractional enrichment at a given time. C. Fractional enrichment for all isotopomers of a-ketoglutarate and succinate in retinas supplied with 5 mM U-^13^C-glucose for 60 minutes. (Error bars indicate S.E.M., n=4). D. Fractional enrichment for all isotopomers of fumarate and succinate in retinas supplied with 5 mM 4-^2^H-glucose for 60 minutes. (Error bars indicate S.E.M., n=3). E. Fractional enrichment for all isotopomers of fumarate and succinate in retinas supplied with 5 mM ^12^C-glucose + 50 μM U-^13^C-malate for 60 minutes. (Error bars indicate S.E.M., n=5).

**SUPPLEMENTAL FIGURE 5.**
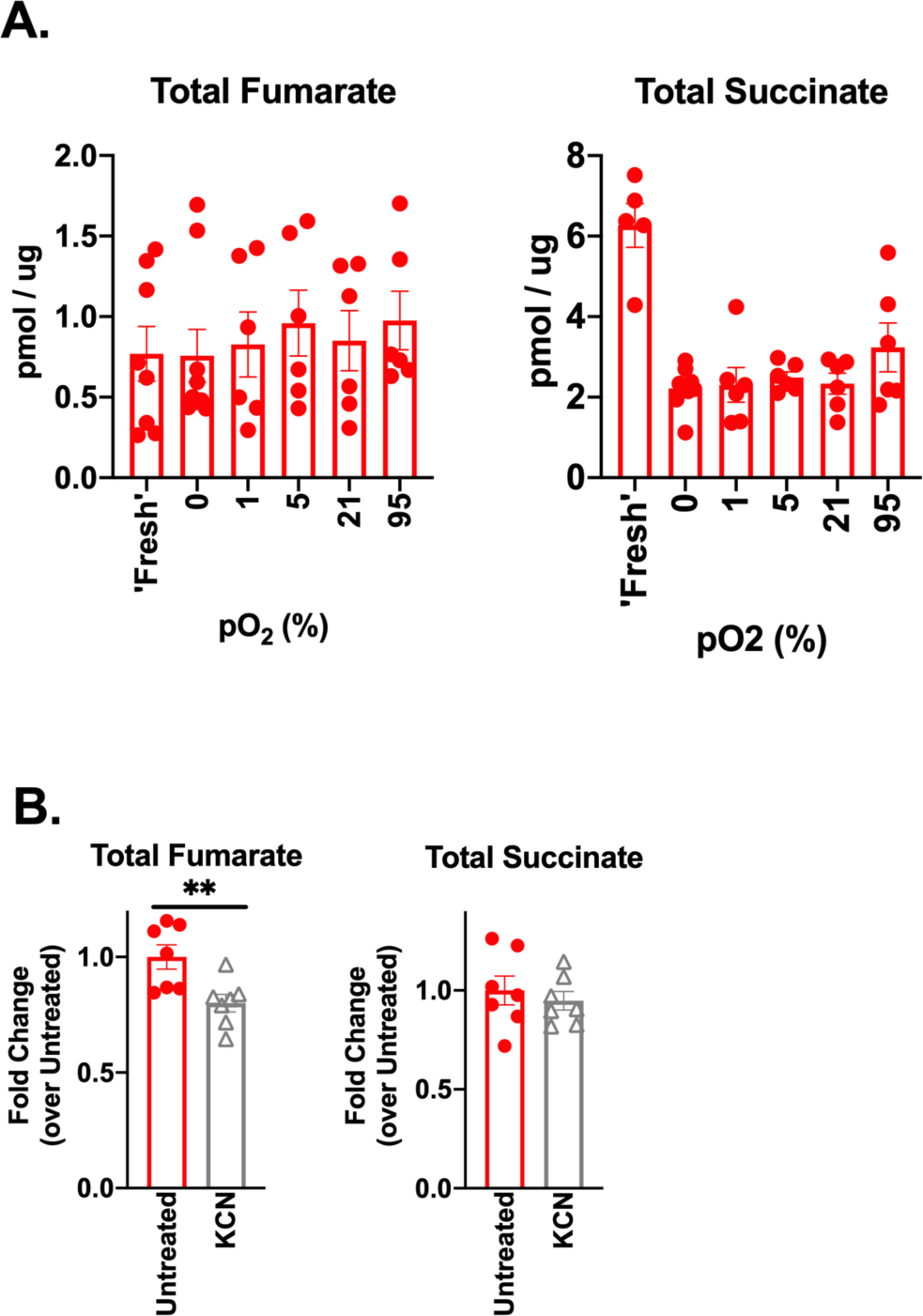
A. Total fumarate and succinate levels in retinas incubated in 5 mM ^12^C-glucose + 50 μM U-^13^C-malate for 5 minutes. “Fresh” indicates that retinas were incubated immediately after dissection. “pO_2_” indicates that retinas were incubated at the specified level of oxygen for 2 hours (in 5 mM ^12^C-glucose) prior to incubation in labeled malate. (Error bars indicate S.E.M., n=6-9). B. Total fumarate and succinate levels in retinas supplied with 5 mM ^12^C-glucose + 50 μM U-^13^C-malate in the presence of KCN. (Error bars indicate S.E.M., n=6).

## MATERIALS AND METHODS

### Immunoblotting

Protein was extracted by homogenizing in RIPA buffer (150 mM NaCl, 1.0% Triton X-100, 0.5% sodium deoxycholate, 0.1% SDS, 50 mM Tris, pH 8.0) and run on 12% polyacrylamide gels. After running, gels were transferred onto PVDF membranes (Millipore, IPFL00010) and blocked for 1 hr at room temperature in LI-COR Odyssey Blocking Buffer (LI-COR, 927-40000). Primary antibodies were diluted in blocking buffer at specified concentrations and incubated overnight at 4°C. Membranes were washed with 1x phosphate buffered saline (PBS) and PBS with 0.1% Tween-20 (PBS-T), then incubated with secondary antibody for 1 hr at 25°C and washed again before imaging. Membranes were imaged and bands were quantified using the LI-COR Odyssey CLx Imaging System (RRID:SCR_014579).

**Table 1.**
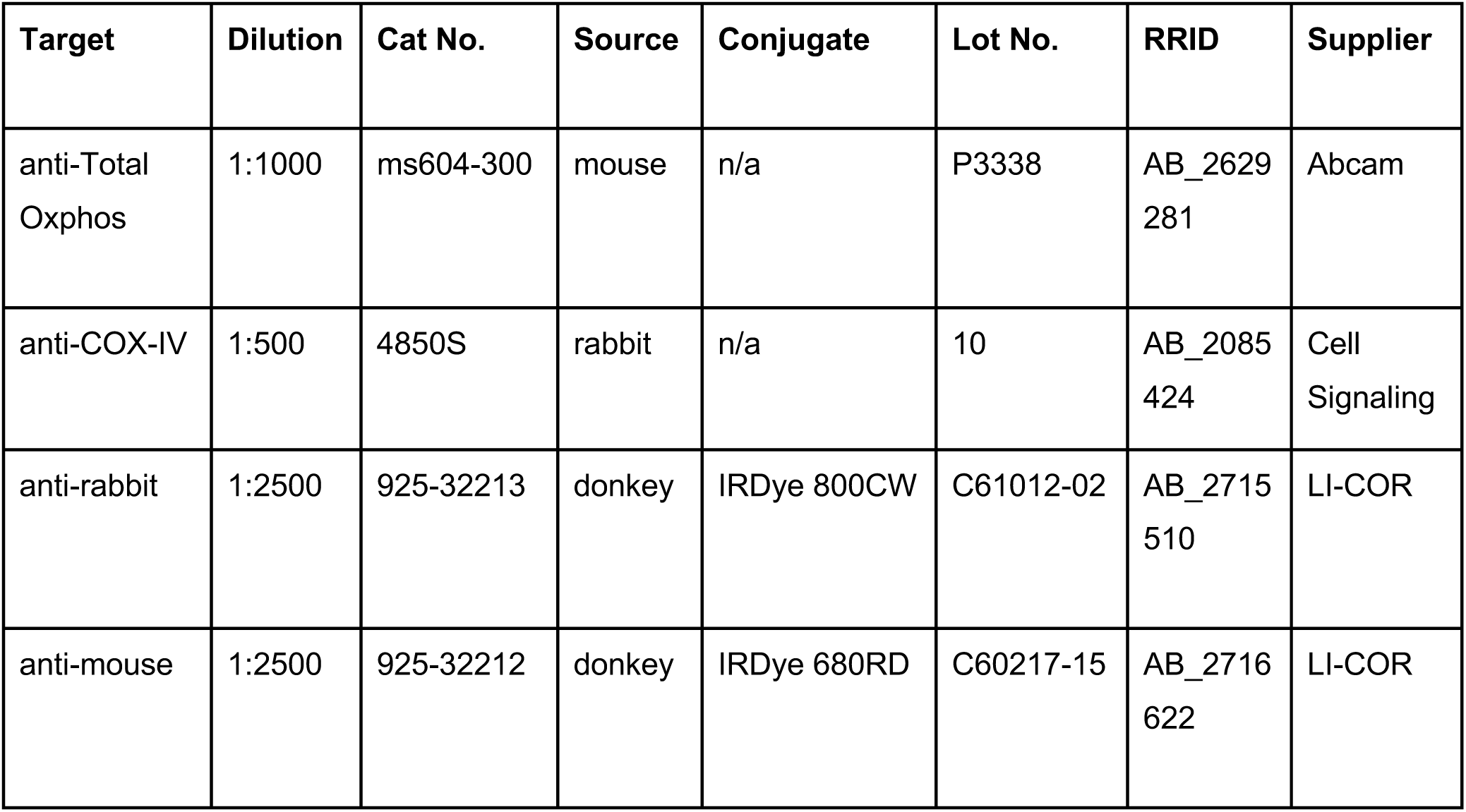
Antibodies for Western Blotting:

| Target | Dilution | Cat No. | Source | Conjugate | Lot No. | RRID | Supplier |
| --- | --- | --- | --- | --- | --- | --- | --- |
| anti-Total Oxphos | 1:1000 | ms604-300 | mouse | n/a | P3338 | AB_2629281 | Abcam |
| anti-COX-IV | 1:500 | 4850S | rabbit | n/a | 10 | AB_2085424 | Cell Signaling |
| anti-rabbit | 1:2500 | 925-32213 | donkey | IRDye 800CW | C61012-02 | AB_2715510 | LI-COR |
| anti-mouse | 1:2500 | 925-32212 | donkey | IRDye 680RD | C60217-15 | AB_2716622 | LI-COR |

### Isotopic Labeling

Krebs-Ringer bicarbonate (KRB) buffer (98.5 mM NaCl, 4.9 mM KCl, 1.2 mM KH_2_PO_4_ 1.2 mM MGSO_4_-7H_2_O, 20 mM HEPES, 2.6 mM CaCl-2H_2_O, 25.9 mM NaHCO_3_) optimized for isotopic labeling experiments in retinas was used in these experiments. Mice were euthanized by awake cervical dislocation and eyes were rapidly enucleated into a dish of Hank’s Buffered Salt Solution (HBSS; Gibco, Cat#: 14025-076). Excess fat and connective tissue was trimmed from each eye, then retinas were dissected away from eyecups and the lens was removed. After dissection, retinas were placed in pre-warmed KRB containing D-[U-^13^C]-glucose (Cambridge Isotope Laboratories, CLM-1396), D-[U-^13^C]-succinate (Cambridge Isotope Laboratories, CLM-1571), D-[U-^13^C]-malate (Cambridge Isotope Laboratories, CLM-8065), or D-[4-^2^H]-glucose, (Omicron Biochemicals, GLC-035), at concentrations specified in each experiment. Retinas were incubated for the specified time points at 37°C (at 5% CO_2_ and room oxygen, unless otherwise specified in the main text), then washed twice in ice-cold PBS and flash frozen in liquid nitrogen.

### Mass Spectrometry Sample Preparation

Metabolites were extracted from both retinas and eyecups using ice-cold 80% MeOH. 150 μL extraction buffer was added to each sample and tissue was disrupted by sonication. Samples were then lyophilized at room-temp until dry. We derivatized extracted metabolites in 10 μL of 20 mg/mL Methoxyamine HCl (Sigma, Cat#: 226904) dissolved in pyridine (Sigma, Cat#: 270970) at 37°C for 90 minutes, and subsequently with 10 μL *tert*-butyldimethylsilyl-N-methyltrifluoroacetamide (Sigma, Cat#: 394882) at 70°C for 90 minutes. Metabolites were analyzed on an Agilent 7890/5975C GC-MS using selected-ion monitoring methods described in previous work.^7–10^ Peaks were manually integrated using MSD ChemStation software (Agilent), and correction for natural isotope abundance was performed using the software Isocor (Millard et al., 2012).

### Curve Fitting

A curve was fit to each time course using Graphpad Prism software assuming a first order reaction by using the equation %*metabolite* = *B* ∗ (1 – *e*^− *kt*^). B is the fractional enrichment at the steady state, k is the rate constant for the reaction, t is time, and %metabolite is the fractional enrichment at a given time.

### Oxygen Consumption Measurement

Oxygen consumption measurements were performed using a perifusion flow-culture system described in **Supplemental Figure 1A** (Sweet et al., 2002, 2004). Retinas and Eyecups were dissected in KRB buffer supplemented with 5 mM glucose, then cut into quarters and loaded into chambers. Quarters from 2 retinas and 4 eyecups were loaded into each chamber and considered one replicate.

